# Estimation of sub-epidemic dynamics by means of Sequential Monte Carlo Approximate Bayesian Computation: an application to the Swiss HIV Cohort Study

**DOI:** 10.1101/085993

**Authors:** Ibeh Neke, Stéphane Aris-Brosou

## Abstract

Our ability to accurately infer transmission patterns of infectious diseases is critical to monitor both their spread and the efficacy of public health policies. The use of phylogenetic methods for the reconstruction of viral ancestral relationships has garnered increasing interest, particularly in the characterization of HIV epidemics and sub-epidemics. In the case of this virus, the Swiss HIV Cohort Study (SHCS) contains a wide breadth of genomic data that have been widely used as a means of applying such methods. However, current approaches for quantifying the epidemiological dynamics of diseases are computationally intensive, and fail to scale well with this magnitude of data. To address this issue, we re-implement an Approximate Bayesian Computation (ABC) approach based on sequential Monte Carlo (SMC). By means of simulations, we demonstrate that our implementation is capable of inferring key epidemiological parameters of the Swiss HIV epidemic accurately, and that sampling intensity has no significant effect on the accuracy of our estimates. Applied to a subset of HIV sequences from the SHCS, we show that we can distinguish sub-epidemics that are circulating in culturally distinct Swiss regions. Given these findings, we propose that ABC-SMC samplers will allow us to evaluate the impact of new public health policies, such as the implementation of a needle exchange program in the case of HIV, based on genetic data sampled before and after the implementation of a new policy.

## Introduction

Understanding the transmission patterns of infectious diseases is critical in order to monitor both their spread and the efficacy of public health policies. This, however, requires quantifying epidemiological dynamics. The most informative quantities are the reproductive ratio (*R*_0_) and the growth rate (*r*) of the epidemic. *R*_0_ represents the number of secondary infections caused by a single infected individual (within a susceptible population) during their entire infectious period (Boskova et al., 2014). Therefore, this statistic informs health officials of the transmission potential in a host population. Generally, *R*_0_ < 1 reflects an epidemic that is under control, while *R*_0_ > 1 reflects the opposite (Li et al., 2011). While the utility of this threshold parameter as a measure of eradication has been questioned (Li et al., 2011), it still provides a critical insight into an epidemic. The parameter *r* represents the speed at which the total number of infected individuals grows in a susceptible population (Volz et al., 2009). So, the magnitudes of *R*_0_ and *r* are used to predict the risk and the extent of spread of a particular outbreak.

Traditionally, both the estimation of these parameters and the reconstruction of transmission events have relied heavily on epidemiological surveys that are valuable, yet time-intensive (Colijn and Gardy, 2014). However, recent developments in the field of phylodynamics have facilitated the use of phylogenetic methods for reconstructing ancestral relationships solely from viral sequence data, thus accelerating the inference of epidemiological parameters (Colijn and Gardy, 2014; Volz et al., 2009). These methods allow us to create pathogen phylogenies that are used to infer critical parameters relating to transmission dynamics within host populations. Two processes that are most commonly used for such inferences are the coalescent and the birth-death model. In essence, the coalescent takes *n* sampled individuals and traces their ancestral lineages backwards in time (Tavaré et al., 1997). Historically, the coalescent has been the standard genealogical model in phylogenetic studies at the population level (Kingman, 1982). However, its inability to account for dense sampling of infected individuals, and its lack of a separate estimation for transmission and death rates, are crucial limitations that make it unsuitable for modeling epidemiological transmission (Boskova et al., 2014).

Conversely, the birth-death model (BDM) provides a stochastic framework for the tree-generating process, thereby overcoming the limitations presented by the coalescent. Unlike the classic coalescent model, the BDM assumes stochastic changes in population size, where an individual can be born or can die at any point in time (Stadler, 2009). In an epidemiological context, a birth event corresponds to an individual becoming infected, while a death event corresponds to the removal of an infected individual from the population (death, treatment, or a change in habits) (Stadler, 2009). The BDM thus makes it possible to estimate parameters such as the transmission rate (λ) and the removal rate (*µ*) of an epidemic, from which both *R*_0_ = λ/*µ* and *r* = λ − *µ* can be derived.

The method most commonly used to infer these parameters relies on Bayesian inference, which enables us to learn something about epidemiological parameters through the estimation of their (joint) posterior distribution(Stadler et al., 2012). This posterior distribution is proportional to the product of the likelihood and of the prior distribution, up to a normalizing constant that, typically, cannot be computed analytically. Numerical estimation of the posterior is then carried out by a Markov chain Monte Carlo (MCMC) sampler that draws from this target distribution. However, the performance of these samplers usually does not scale very well with increasingly large data sets (Aris-Brosou and Rodrigue, 2012). Furthermore, in cases where the likelihood function is too complex to derive (Rayner and MacGillivray, 2002; Yildirim et al., 2015), an alternative approach is required. Hence, Approximate Bayesian Computation (ABC) has become a central framework for parameter inference (Toni et al., 2009). ABC methods circumvent the issue of costly MCMC calculations by resorting to simulations, which rely on a parametric model: at each step of the algorithm, a vector of parameters is drawn from a prior distribution, and used to simulate data. Both observed and simulated data are then condensed into a single value (either a real number or a vector) by means of a summary statistic. The vector of parameters used at this step is kept for inference (stored) if the distance between the summary statistics of the observed and simulated data is less than a tolerance value (*ϵ*). In the simplest ABC algorithm, a new vector of parameters is drawn at random at the next step. This process eventually samples from an approximation of the posterior distribution of the parameters of interest (Pritchard et al., 1999; Beaumont et al., 2002).

One of the major limitations of this naïve ABC sampler is that the acceptance rate is low when the prior distribution strongly differs from the posterior distribution. To address this issue, an MCMC-based ABC sampler was introduced, where the move from a current state to the next is made by modifying the current state (Marjoram et al., 2003). Although ABC-MCMC is guaranteed to converge to the target approximate posterior distribution, it also possesses some potential disadvantages. Notably, the efficiency of the algorithm is relatively low due to the correlated nature of samples coupled with a potentially low acceptance rate (Toni et al., 2009). In order to alleviate this issue, a sequential Monte Carlo (SMC)-based sampler was described. By propagating sampled parameter values (particles) through a sequence of intermediate distributions defined by a tolerance schedule, parameters that represent a sample from the target posterior distribution are eventually obtained (Toni et al., 2009).

The success of this approach is a function of the ease with which simulations are performed, the distance metric that is used, and more critically, the summary statistic that is employed (Marjoram et al., 2003). To compute the distance between a simulated and an observed phylogeny, each data set must be summarized in a manner that captures the information contained therein. Various summary statistics have been proposed for characterizing phylogenetic trees, such as Sackin’s index (Sackin, 1972; Blum and Francois, 2005) or the total cophenetic index (Mir et al., 2013), but very few of these can effectively (*sufficiently*) capture the underlying epidemiological dynamics of a viral phylogeny. Often, however, the comparison of two phylogenies has been made possible through the use of their Lineages-Through-Time (LTT) curves (Nee et al., 1994; Paradis, 2011). Since the LTT curve of a phylogenetic tree depicts the number of lineages in the phylogeny through time, it reveals the distinct patterns of speciation and extinction that are specific to each tree. To avoid issues that arise when comparing trees of different sizes, Janzen et al. (2015) proposed the normalized LTT (nLTT) to compare trees with differing height and number of extant tips.

Although attempts at applying ABC methods to the parameter inference problem have been made, these approaches are computationally intensive, employ poorly-informative summary statistics, and fail to scale well with large data sets (Leventhal et al., 2012; Höhna et al., 2011; Stadler et al., 2012). Here, we test a new implementation of an ABC-SMC sampler to infer epidemiological parameters, based on the nLTT. By means of simulations, we show that ABC-SMC yields reliable and accurate estimates of the transmission (λ) and removal (*µ*) rates of an epidemic. We validate our approach using data from the Swiss HIV-1 epidemic as a case study (Cohort profile, 2010), where genetic data obtained from two different regions of the country are expected to exhibit distinct transmission patterns. We report specific parameter estimates for each region, irrespective of the subsampling of these data, highlighting the sensitivity of our approach.

## Materials and Methods

Taking inspiration from a recent implementation of the birth-death model (Stadler, 2010; Stadler et al., 2012), our epidemiological model follows a constant-rate BDM with incomplete species sampling (*ρ*) (see also Stadler, 2009; Yang and Rannala, 1997; Nee et al., 1994). Through this forward-in-time description of the epidemiological process, the BDM specifies sampling intensity and thus enables explicit simulation of the serially sampled infected class in our ABC approach. While the original BDM implementation in R (package TreeSim) enables the inference of epidemiological dynamics under constant or episodic processes (Stadler, 2011), its utility for such inference is limited to relatively small phylogenetic trees. A more recently developed BDM R-package, called TESS (Höhna et al., 2015), provides the same features offered in TreeSim, but with a more flexible framework for specifying diversification models. As such, TESS was used here as a means of simulating phylogenetic trees – for for the purpose of inference (by ABC-SMC) and of performance assessment (by simulations).

### ABC-SMC estimation under the BDM

To infer the epidemiological parameters imprinted on each phylogenetic tree, we employed an ABC-SMC method (Toni et al., 2009). In ABC-SMC, a series of parameter values (often termed particles) are first sampled from the prior distribution. These particles are then propagated through a sequence of intermediate distributions defined by a tolerance schedule, where the threshold value (*ϵ*) for acceptance is sequentially lowered. This threshold value is based on a summary statistic which captures the information represented by the data. As the acceptance threshold approaches zero for each sequential population of particles, the intermediate distributions gradually morph towards the target posterior distribution. This population approach alleviates the major issue of getting stuck in areas of low probability, usually faced with the more standard ABC-MCMC approach (Toni et al., 2009).

The ABC-SMC approach presented here was based on the nLTT R package developed by Janzen et al. (2015). However, modifications were made to the original source code (version 1.1, accessed December 7, 2015) to increase functionality and range of applicability. Specifically, the prior for the density function was changed from a normal prior to a uniform prior on [0, 5], so that it is consistent with the uniform prior used to generate the birth-death parameters (λ and *µ*). Such a prior also covers the most realistic values for these parameters (Janzen et al., 2015), and ensures that neither parameter becomes too large. Additionally, the λ and *µ* prior functions were adjusted so that the tree simulating process would not be compromised when *µ* > λ. We found that these modifications significantly improved the functionality of the ABC-SMC code provided by Janzen and colleagues.

The SMC *ϵ* schedule was set following the procedure described in Janzen et al. (2015) and Del Moral et al. (2012). Briefly, the schedule is decreased based on an exponential progression, such that the *ϵ* value at time *t* (current step in the sequence) is *ϵ*_*t*_ = *ϵ*_0_ exp(−0.5*t*), starting from an *ϵ*_0_ = 0.2. For each iteration of the SMC algorithm, 1000 particles were used, and convergence was assumed when the acceptance rate of the proposed particles dropped below 0.001.

Although the performance of ABC-SMC samplers has historically been superior to that of the basic rejection sampler and ABC-MCMC (Lenormand et al., 2013; Toni et al., 2009), the efficiency of the ABC-SMC algorithm hinges on the choice of an appropriate summary statistic for the data. Given that the data observed here are in the form of phylogenetic trees, we hypothesized that the normalized LTT (nLTT) curves of each tree would serve as appropriate summary statistics. Using these curves, two phylogenies can effectively be summarized and compared against one another. We therefore implemented this summary statistic into our ABC-SMC algorithm as a means of comparing our simulated and observed phylogenies. We calculated the nLTT statistic using the function nLTTstat from the R-package nLTT in R 3.2.1 (Janzen et al., 2015).

### Performance assessment

We simulated birth-death trees under several epidemiological scenarios in order to assess the ability of our method to infer both removal (*µ*) and transmission (λ) rates accurately. The parameters used in this model were λ, *µ, ρ* (sampling probability), T (total number of extant tips), and epidemic age. The possible empirical values for λ and *µ* were based on a geometric sequence from 0 to 5 with common ratio 1.5, yielding values {0.5, 0.75, 1.125, 1.6875, 2.53125, 3.796875} for λ and {0.15, 0.225, 0.3375, 0.50625, 0.759375, 1.1390625} for *µ*. The parameter *µ* was restricted as to be smaller than λ in order to avoid simulating uninformative phylogenies. Trees were simulated with sampling probabilities (*ρ*) of 1, 0.5, and 0.25. The number of taxa (T) was set to 500, 1000, 10,000, and 20,000. Age was also varied, with values set to 10, 20, and 30. These chosen values for number of taxa and epidemic age represent a wide range of potential real-world epidemic scenarios. This factorial design represents a total of 1080 simulation conditions. Due to the size of this simulation study, we only performed 25 replicates for each simulation condition. The accuracy of our algorithm was measured using the mean squared error (MSE).

### Application to the Swiss HIV-1 sequence data

We used HIV-1 polymerase sequence data from the Swiss HIV Cohort Study (Cohort profile, 2010). The available Swiss HIV pol sequences (570) (Kouyos et al., 2010) were aligned using MUSCLE version 3.8.31 (Edgar, 2004) and manually inspected using JalView (Waterhouse et al., 2009). The sequences used were divided based on the presence of a large indel in one group of sequences: one sequencing center starts at the end the protease, at nucleotide 297, while the other one starts from within the reverse transcriptase, at nucleotide 84. This phylogeographic pattern therefore reflects two groups of sequences representing two culturally unique regions within Switzerland that were sampled. Since the identity of each group could not be inferred from GenBank annotations, we labeled them as Region 1 (252 sequences) and Region 2 (318 sequences), respectively.

In order to determine the age of each tree (length of the epidemic), each alignment was analyzed using a relaxed molecular clock (Drummond et al., 2006) as implemented in BEAST v1.8.0 (Drummond and Rambaut, 2007) using a GTR+Γ_4_ model of nucleotide substitution (Lanave et al., 1984; Tavaré, 1986; Yang, 1996; Aris-Brosou and Rodrigue, 2012), assuming an uncorrelated lognormal relaxed clock and a constant-population coalescent prior. We ran two separate MCMC samplers for each alignment (10^10^ states each) to check for convergence, visually assessed these with Tracer (tree.bio.ed.ac.uk/software/tracer), and conservatively removed 10% of the states as burn-in.

Using the two trees obtained through the relaxed molecular clock analyses, we applied our ABC-SMC algorithm to the Swiss HIV data. The empirical estimates for parameters *µ* and λ were obtained for Regions 1 and 2. As the sequence data deposited in GenBank (252 + 318 sequences) is a subset of the data used in Kouyos et al. (2010) (*n* = 5700), we tested whether further subsetting of the data would return consistent parameter estimates. For this, we split each alignment into two sub-alignments, and ran the ABC-SMC algorithm again. This procedure was conducted ten times, yielding twenty sub-alignments. All the scripts and data we used have been deposited at github.com/sarisbro.

## Results and Discussion

To alleviate the computational burden incurred by rigorous epidemiological modeling and MCMC sampling, we re-implemented an epidemiological model employing an ABC-SMC sampler. We first validated our approach by means of an extensive simulation study, and then tested our ability to distinguish sub-epidemics using data from the SHCS.

### Simulated data

Based on 25 replicates for each parameter setting, we assessed the ability of our method to recover the values for each combination of λ and *µ*. Simulation results show that the SMC algorithm converges upon the true posterior within 11 populations of the ABC-SMC sampler (Figure 1), using solely the nLTT as a summary statistic.

**Figure 1.**
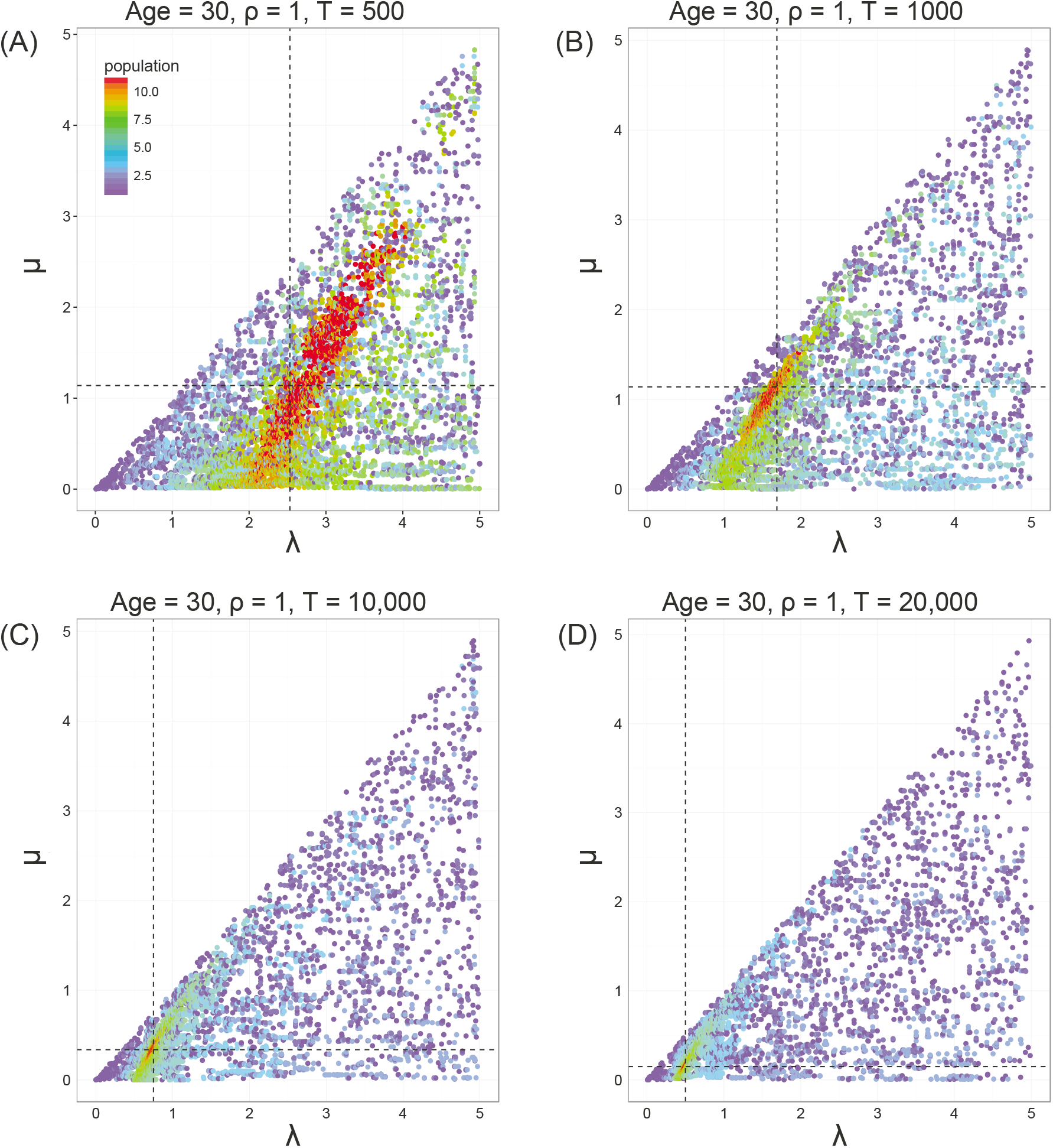
ABC-SMC estimates of the posterior distribution for four taxa settings. Number of populations required for convergence (max = 11) is indicated using color spectrum. Estimated parameter values for each setting are indicated with dashed lines. (A): (λ=2.531, *µ*=1.139). (B): (λ=1.125, *µ*=1.139). (C): (λ=0.750, *µ*=0.338). (D): (λ=0.500, *µ*=0.150). Particles per iteration = 1000. N = 25 replicate trees.

Such quick convergence supports our hypothesis that the nLTT is an appropriate and informative summary statistic, given the nature of our data. While previous authors (Janzen et al., 2015) compared the performance of the nLTT to other phylogenetic summary statistics within an ABC-SMC framework, the parameter space they explored was limited. In particular, birth-death trees were only simulated under age 10 and no more than 178 tips (extant taxa). Here, we cover a broader parameter space, presenting a greater number of epidemiological scenarios. Still, the quick convergence onto the true posterior distribution further demonstrates that the nLTT is a suitable summary statistic for parameter inference (Janzen et al., 2015).

The marginal MSEs for λ and *µ* (Figure 2) illustrate that our SMC approach was successful in recovering the real values of these parameters using only the nLTT of each tree as a summary statistic. Marginal MSEs for λ were surprisingly low, with an upper bound of 0.2 for all combinations of age, number of taxa, and sampling intensity. While the marginal MSEs obtained for *µ* were also reasonably low, our method was not as accurate in recovering these estimates as it was for λ. This is evident when comparing the upper bound of marginal MSEs for *µ* (0.5) versus λ (0.2).

**Figure 2.**
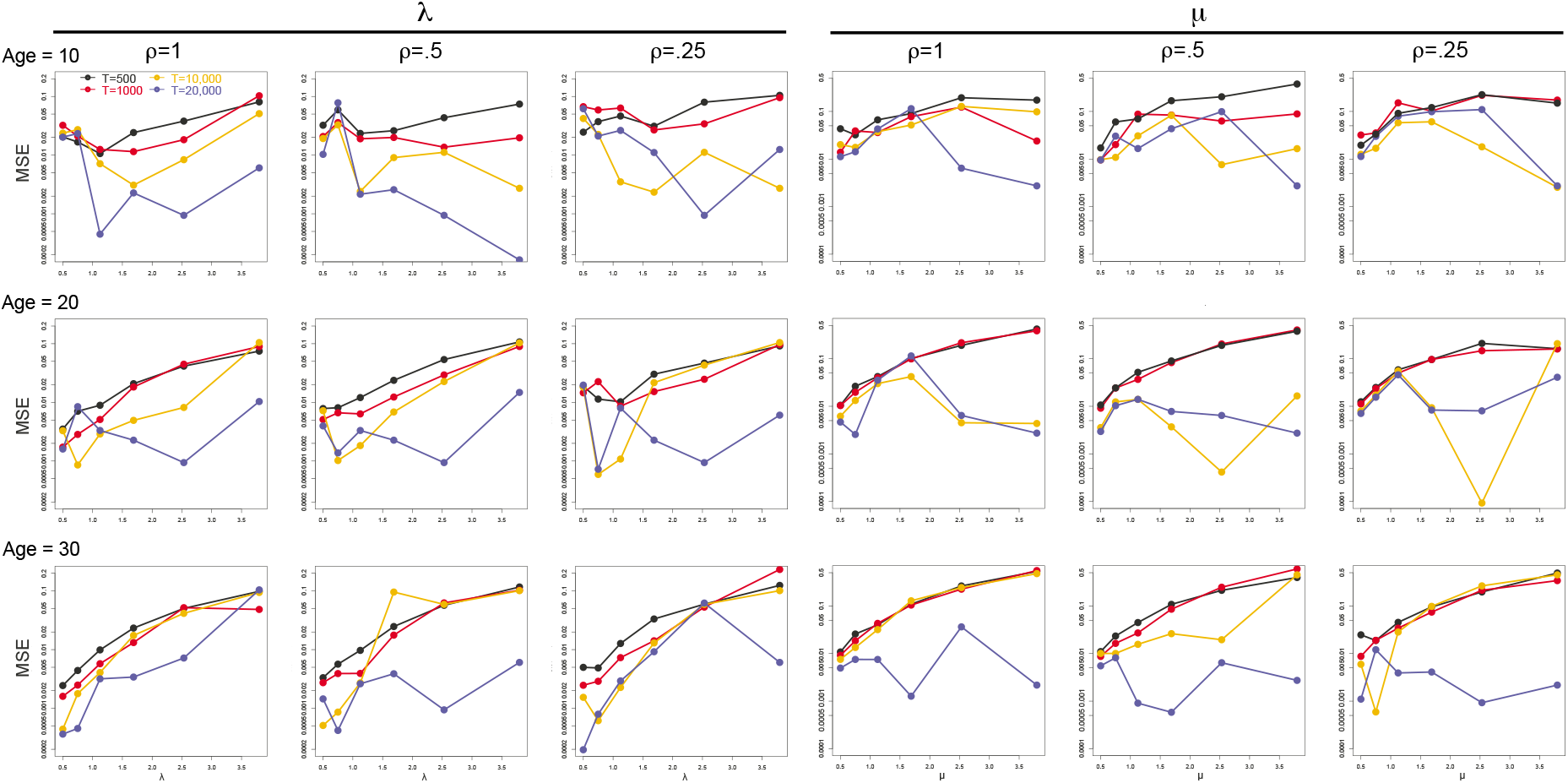
Marginal MSEs for *λ* and *µ*. Mean squared errors for all epidemiological settings are shown. Y-axis is log_10_-transformed. Each point represents an MSE value over 25 replicate trees.

Simulations therefore show that the transmission rate is more accurately estimated than the removal rate. Inferring the removal rate seems to be complicated by the non-independence between λ and *µ* (Figure 1). In our approach, we limited *µ* to values less than λ to avoid the high computational cost of trees simulated with a low birth rate and high death rate. This constraint is potentially limiting our ability to accurately infer *µ*, as λ might leave a more detectable imprint than *µ* on each reconstructed tree, thereby increasing the estimated growth rate (*r*). Consequently, certain regions of the parameter space correspond to very high estimation accuracy (low MSEs), while other regions give rise to low accuracy (high MSEs; Figure 3), as previously reported (Capaldi et al., 2012). This issue is further compounded by the inevitable epidemiological correlation between the birth rate and death rate. Previous work (Capaldi et al., 2012) has shown that the transmission and removal rates in an epidemiological model are highly correlated, with the magnitude of the correlation efficient being strongly dependent on the value of *R*_0_. As *R*_0_ approaches 1, the correlation coefficient peaks, and standard errors for point estimates increase significantly. Such a pattern can be observed in Figure 3. This correlation thus contributes to the identifiability constraints we observed here.

**Figure 3.**
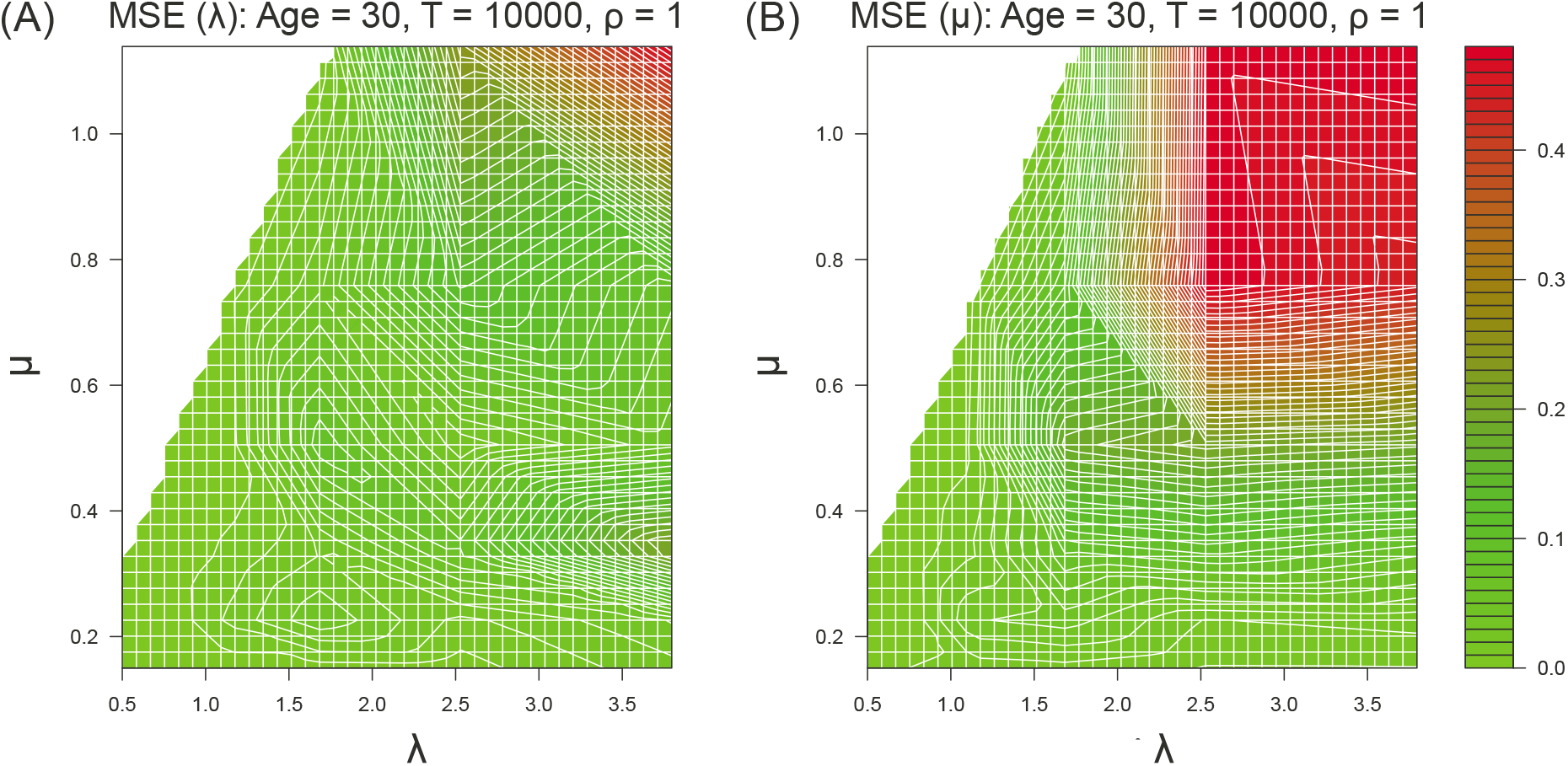
Joint MSE heatmaps. Mean squared error for a set age, taxa size, and sampling probability for λ (A) and *µ* (B). Axes show range of simulated values for all trees (λ ≤ *µ*).

Although the correlation between these two parameters has been examined in previous studies (Capaldi et al., 2012; Li and Vu, 2013; Wu et al., 2008), there is still no consensus on how identifiability constraints can be overcome. It has been noted that the time at which observations are made during an epidemic has a significant bearing on the precision of the estimates (Capaldi et al., 2012). Sensitivity functions have been proposed as a means of determining points in time when the epidemiological system is most sensitive to parameter fluctuations (Capaldi et al., 2012; Banks et al., 2007b,a). Analysis based on such functions illustrates that the correlation between estimates of λ and *µ* is highest during the exponential growth phase, early in the epidemic (Capaldi et al., 2012). As the constant-rate BDM employed here is limited to the exponential phase of an epidemic, such a strong correlation between λ and *µ* is expected. Therefore, this BDM technique can be further developed in future studies so that it also encompasses the saturation phase of an epidemic.

Our results also suggest that accuracy is solely affected by the number of taxa on the simulated tree. Essentially, as the number of taxa increases, our ability to accurately infer these epidemiological parameters increases as well (Figure 2). Accuracy is particularly high for trees simulated with 1000 or more sequences (Figure 1). Recent studies have also noted that parameter inference using the BDM is biased towards larger population sizes (Stadler et al., 2012; Boskova et al., 2014). This suggests that larger phylogenies provide much more information about the dynamics of an epidemic, thereby facilitating the process of parameter inference. Uniquely here, we also find that sampling intensity (*ρ*) and age bear no statistically significant impact on accuracy (Figure 4). These results suggest that our approach is robust to most sampling designs (sampling effort and/or genetic diversity) and epidemic ages.

**Figure 4.**
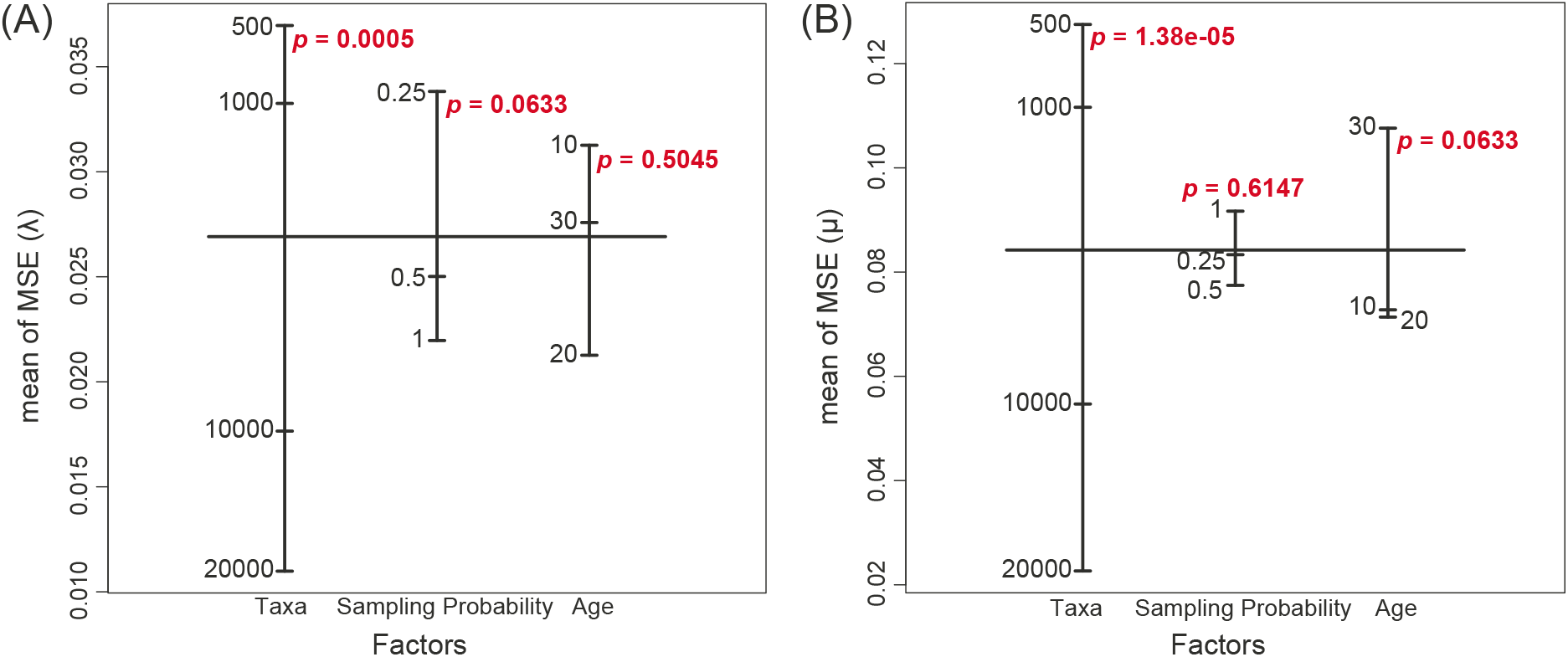
Three-factorial ANOVA for marginal MSEs. Three-factorial ANOVA for marginal MSEs. ANOVA results for λ (A) and *µ* (B). Independent factors listed are number of taxa, sampling probability, and age. P-value for each factor is denoted in red.

### Swiss HIV-1 parameter estimates

We applied our method to HIV-1 sequence data from the Swiss HIV Cohort Study (Cohort profile, 2010). Using the two dated phylogenetic trees obtained from the relaxed clock analyses, we inferred the transmission and removal rates for Regions 1 and 2. We tested whether our method was capable of distinguishing between the transmission dynamics of these two geographically similar but culturally distinct samples.

While the trees corresponding to each region (Figure 5A-B) suggested that the two phylogenies were very similar with respect to topology, our algorithm recovered two distinct epidemiological patterns (Figure 5C). The contour plots represent the joint posterior distribution of λ and *µ* approximated by the ABC-SMC algorithm for the two regions. Subsequent analysis of the subset alignments confirmed these estimations, as the posterior distributions for all replicates overlapped within each region, but not between regions (Figure 6). The parameter estimates and 95% HPD intervals for each region are listed in Table 1. Estimates using the upper and lower bounds of the 95% HPD intervals for age are listed in Table 2 for reference.

**Figure 5.**
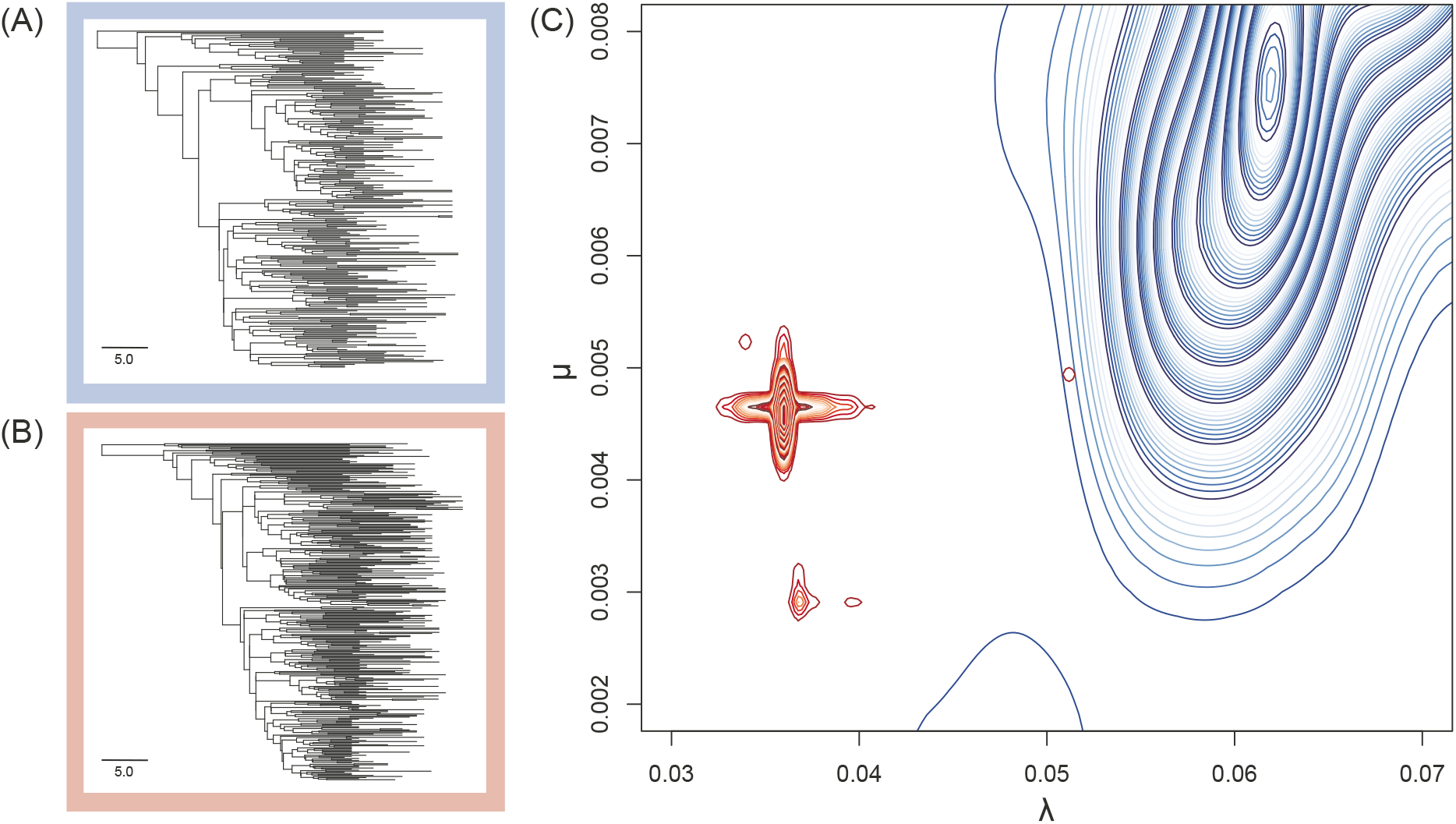
Swiss HIV-1 regional trees and parameter estimates. (A-B): Dated phylogenetic BEAST trees for Swiss HIV sub-clusters under a relaxed molecular clock. Scale is in years. (C): Joint posterior distribution of λ and *µ* approximated with the ABC-SMC algorithm. Distribution for Region 1 is shown in blue, and Region 2 is in red.

**Figure 6.**
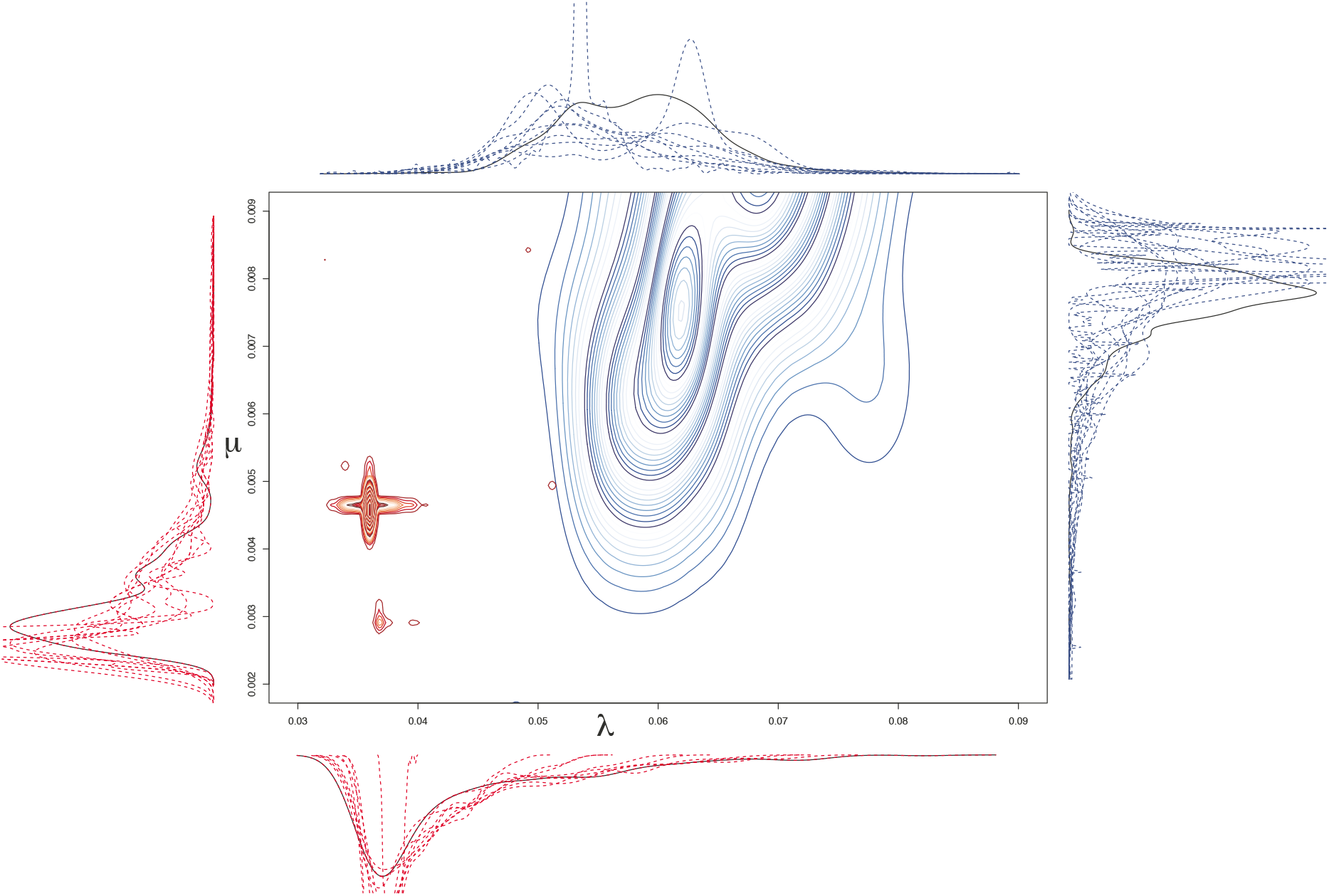
Swiss HIV-1 parameter estimates along with subset analyses. Joint posterior distribution of λ and *µ* approximated with the ABC-SMC algorithm. Distribution for Region 1 is shown in blue, and Region 2 is in red. Marginal posterior distributions of each region are shown in corresponding colors on each axis for the complete alignments (solid lines) and for the subset alignments (20; dotted lines).

**Table 1.**
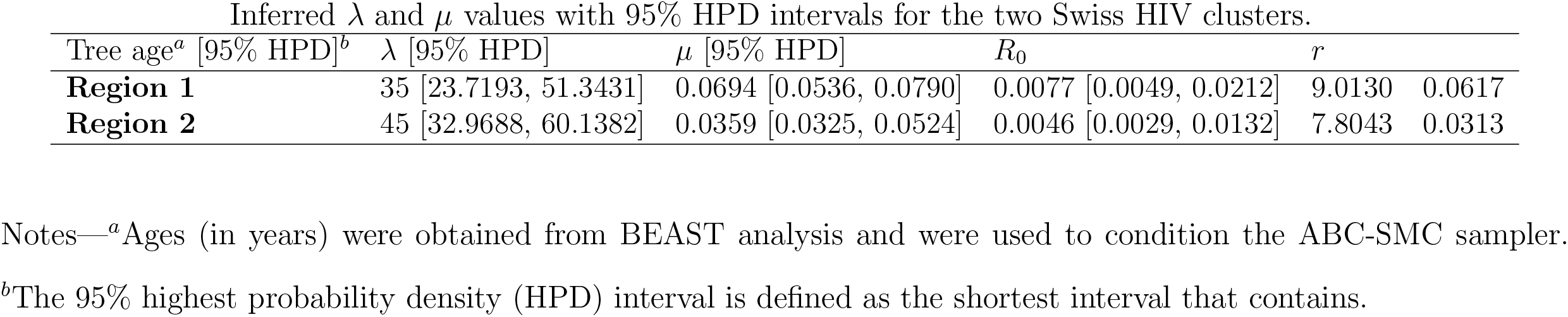
Inferred *λ* and *µ* values with 95% HPD intervals for the two Swiss HIV clusters. *R*_0_ and *r* values were obtained using λ and *µ* estimates.

| Inferred $\lambda$ and $\mu$ values with 95% HPD intervals for the two Swiss HIV clusters. | | | | | |
| --- | --- | --- | --- | --- | --- |
| Tree age<sup>a</sup> [95% HPD]<sup>b</sup> | $\lambda$ [95% HPD] | $\mu$ [95% HPD] | $R_0$ | $r$ | |
| **Region 1** | 35 [23.7193, 51.3431] | 0.0694 [0.0536, 0.0790] | 0.0077 [0.0049, 0.0212] | 9.0130 | 0.0617 |
| **Region 2** | 45 [32.9688, 60.1382] | 0.0359 [0.0325, 0.0524] | 0.0046 [0.0029, 0.0132] | 7.8043 | 0.0313 |

Notes—<sup>a</sup>Ages (in years) were obtained from BEAST analysis and were used to condition the ABC-SMC sampler.
<sup>b</sup>The 95% highest probability density (HPD) interval is defined as the shortest interval that contains.

**Table 2.** Inferred *λ* and *µ* values using the upper and lower HPD bounds for age. *R*_0_ and *r* values were obtained using λ and *µ* estimates. Inferred λ and *µ* values using the upper and lower HPD bounds for age.

| Tree age | $\lambda$ [95% HPD] | $\mu$ [95% HPD] | $R_0$ | $r$ |
| --- | --- | --- | --- | --- |
| **Region 1** | 23 | 0.0322 [0.0266, 0.0570] | 0.0157 [0.0091, 0.0316] | 2.0510 0.0165 |
|  | 51 | 0.0046 [0.0027, 0.0075] | 0.0011 [0.0002, 0.0029] | 4.1818 0.0035 |
| **Region 2** | 32 | 0.0058 [0.0031, 0.0069] | 0.0007 [0.0006, 0.0009] | 8.2857 0.0051 |
|  | 60 | 0.0754 [0.0636, 0.0855] | 0.0099 [0.0081, 0.0296] | 7.6162 0.0655 |

Our results indicate that during the Swiss HIV epidemic, Region 1 had a higher growth rate than Region 2, suggesting that the pathogen was spreading at a faster pace throughout the population. Moreover, Region 1 had a higher basic reproductive ratio than Region 2 (Table 1). The *R*_0_ estimates for Regions 1 and 2 are significantly greater than 1, indicating that the probability of infection and the extent of spread of the disease within these regions in Switzerland were high. While this is in line with previous predictions based on samples from the Swiss HIV Cohort Study (Stadler et al., 2012), no study to date has conducted any analysis on the specific sequences and epidemic time-frame used here, so comparisons cannot be made.

With no available data connecting the sequences to their exact origins, no conclusions can be made about the cultural effects of each region on the above estimates. Nonetheless, one factor which may have influenced the different epidemiological patterns in each region is their respective composition according to transmission group. The subsamples studied here may be dominated by different risk groups (i.e. men having sex with men, heterosexuals, and intravenous drug users) (Kouyos et al., 2010; Stadler et al., 2012), which may have an impact on the underlying dynamics of the subepidemics.

### Conclusions

Various studies have attempted to estimate epidemiological parameters directly from viral sequence data. Past approaches based on the coalescent as a transmission model were unable to directly infer transmission and death rates, and they had to resort to a sophisticated independent estimate for the duration of infectiousness (Volz et al., 2009; Frost and Volz, 2010). More recently, Stadler *et al.* (Stadler et al., 2012) applied a Bayesian BDM to the parameter estimation problem. While they accurately inferred *R*_0_ values, their approach was incapable of accurately estimating the transmission and death rates separately. Here, we show that our method can accurately recover the transmission and removal rates of simulated empirical trees. Most importantly, our reanalysis of data from the Swiss HIV Cohort Study shows that we are able to distinguish transmission patterns in two culturally different parts of Switzerland.

Our ability to infer the epidemiological parameters of two subsamples with very similar phylogenetic structures, highlights the utility and power of our method. Our approach provides useful information about an outbreak even when epidemiological data is incomplete, and it is thus informative within the earliest points of an outbreak. In light of these results, we propose that our method will make it possible for future studies to evaluate the impact of new public health policies (e.g., implementation of a needle exchange program in the case of HIV) based on genetic data sampled before and after the implementation of a new policy.

## Acknowledgements

We thank Thijs Janzen for discussions about the source code of the nLTT package, and Roger Kouyos for sharing some information about the HIV data used in this study. This work was supported by the Natural Sciences Research Council of Canada and by the Canada Foundation for Innovation (S.A.B.) and by the University of Ottawa (N.I.).

## References

Aris-Brosou, S., Rodrigue, N., 2012. The essentials of computational molecular evolution. Methods Mol Biol 855, 111–52. doi:10.1007/978-1-61779-582-4_4.

Banks, H., Dediu, S., Ernstberger, S.L., 2007a. Sensitivity functions and their uses in inverse problems. Journal of Inverse and Ill-posed Problems jiip 15, 683–708.

Banks, H., Ernstberger, S.L., Grove, S.L., 2007b. Standard errors and confidence intervals in inverse problems: sensitivity and associated pitfalls. Journal of Inverse and Ill-posed Problems jiip 15, 1–18.

Beaumont, M.A., Zhang, W., Balding, D.J., 2002. Approximate Bayesian computation in population genetics. Genetics 162, 2025–2035.

Blum, M.G., Francois, O., 2005. On statistical tests of phylogenetic tree imbalance: the Sackin and other indices revisited. Mathematical Biosciences 195, 141–153.

Boskova, V., Bonhoeffer, S., Stadler, T., 2014. Inference of epidemiological dynamics based on simulated phylogenies using birth-death and coalescent models. PLOS Comput Biol 10, e1003913.

Capaldi, A., Behrend, S., Berman, B., Smith, J., Wright, J., Lloyd, A.L., 2012. Parameter estimation and uncertainty quantication for an epidemic model. Mathematical Biosciences and Engineering, 553.

Cohort profile, 2010. Cohort profile: the Swiss HIV Cohort study. International Journal of Epidemiology 39, 1179–1189.

Colijn, C., Gardy, J., 2014. Phylogenetic tree shapes resolve disease transmission patterns. Evolution, Medicine, and Public Health 2014, 96–108.

Del Moral, P., Doucet, A., Jasra, A., 2012. An adaptive sequential Monte Carlo method for approximate Bayesian computation. Statistics and Computing 22, 1009–1020.

Drummond, A.J., Ho, S.Y., Phillips, M.J., Rambaut, A., 2006. Relaxed phylogenetics and dating with confidence. PLoS Biol 4, e88.

Drummond, A.J., Rambaut, A., 2007. Beast: Bayesian evolutionary analysis by sampling trees. BMC Evolutionary Biology 7, 214.

Edgar, R.C., 2004. Muscle: multiple sequence alignment with high accuracy and high throughput. Nucleic Acids Research 32, 1792–1797.

Frost, S.D., Volz, E.M., 2010. Viral phylodynamics and the search for an ‘effective number of infections’. Philosophical Transactions of the Royal Society of London B: Biological Sciences 365, 1879–1890.

Höhna, S., May, M.R., Moore, B.R., 2015. TESS: an R package for efficiently simulating phylogenetic trees and performing Bayesian inference of lineage diversification rates. Bioinformatics, btv651.

Höhna, S., Stadler, T., Ronquist, F., Britton, T., 2011. Inferring speciation and extinction rates under different sampling schemes. Molecular Biology and Evolution 28, 2577–2589.

Janzen, T., Höohna, S., Etienne, R.S., 2015. Approximate Bayesian computation of diversification rates from molecular phylogenies: introducing a new efficient summary statistic, the nltt. Methods in Ecology and Evolution 6, 566–575.

Kingman, J.F.C., 1982. The coalescent. Stochastic Processes and their Applications 13, 235–248.

Kouyos, R.D., Von Wyl, V., Yerly, S., Böni, J., Taffé, P., Shah, C., Börgisser, P., Klimkait, T., Weber, R., Hirschel, B., et al., 2010. Molecular epidemiology reveals long-term changes in HIV type 1 subtype B transmission in Switzerland. Journal of Infectious Diseases 201, 1488–1497.

Lanave, C., Preparata, G., Sacone, C., Serio, G., 1984. A new method for calculating evolutionary substitution rates. Journal of Molecular Evolution 20, 86–93.

Lenormand, M., Jabot, F., Deffuant, G., 2013. Adaptive approximate Bayesian computation for complex models. Computational Statistics 28, 2777–2796.

Leventhal, G.E., Kouyos, R., Stadler, T., Von Wyl, V., Yerly, S., Böni, J., Cellerai, C., Klimkait, T., Günthard, H.F., Bonhoeffer, S., 2012. Inferring epidemic contact structure from phylogenetic trees. PLoS Comput Biol 8, e1002413.

Li, J., Blakeley, D., et al., 2011. The failure of *r*_0_. Computational and mathematical methods in medicine 2011.

Li, P., Vu, Q.D., 2013. Identification of parameter correlations for parameter estimation in dynamic biological models. BMC Systems Biology 7, 91.

Marjoram, P., Molitor, J., Plagnol, V., Tavaré, S., 2003. Markov chain Monte Carlo without likelihoods. Proceedings of the National Academy of Sciences 100, 15324–15328.

Mir, A., Rosselló, F., et al., 2013. A new balance index for phylogenetic trees. Mathematical Biosciences 241, 125–136.

Nee, S., May, R.M., Harvey, P.H., 1994. The reconstructed evolutionary process. Philosophical Transactions of the Royal Society of London B: Biological Sciences 344, 305–311.

Paradis, E., 2011. Time-dependent speciation and extinction from phylogenies: A least squares approach. Evolution 65, 661–672.

Pritchard, J.K., Seielstad, M.T., Perez-Lezaun, A., Feldman, M.W., 1999. Population growth of human y chromosomes: a study of y chromosome microsatellites. Molecular Biology and Evolution 16, 1791–1798.

Rayner, G.D., MacGillivray, H.L., 2002. Numerical maximum likelihood estimation for the g-and-k and generalized g-and-h distributions. Statistics and Computing 12, 57–75.

Sackin, M., 1972. “good” and “bad” phenograms. Systematic Biology 21, 225–226.

Stadler, T., 2009. On incomplete sampling under birth-death models and connections to the sampling-based coalescent. Journal of Theoretical Biology 261, 58–66.

Stadler, T., 2010. Sampling-through-time in birth-death trees. Journal of Theoretical Biology 267, 396–404.

Stadler, T., 2011. Simulating trees with a fixed number of extant species. Systematic Biology, syr029.

Stadler, T., Kouyos, R., von Wyl, V., Yerly, S., Böoni, J., Bürgisser, P., Klimkait, T., Joos, B., Rieder, P., Xie, D., et al., 2012. Estimating the basic reproductive number from viral sequence data. Molecular Biology and Evolution 29, 347–357.

Tavaré, S., 1986. Some probabilistic and statistical problems in the analysis of dna sequences. Lectures on mathematics in the life sciences 17, 57–86.

Tavaré, S., Balding, D.J., Griffiths, R.C., Donnelly, P., 1997. Inferring coalescence times from DNA sequence data. Genetics 145, 505–518.

Toni, T., Welch, D., Strelkowa, N., Ipsen, A., Stumpf, M.P., 2009. Approximate Bayesian computation scheme for parameter inference and model selection in dynamical systems. Journal of the Royal Society Interface 6, 187–202.

Volz, E.M., Pond, S.L.K., Ward, M.J., Brown, A.J.L., Frost, S.D., 2009. Phylodynamics of infectious disease epidemics. Genetics 183, 1421–1430.

Waterhouse, A.M., Procter, J.B., Martin, D.M., Clamp, M., Barton, G.J., 2009. Jalview version 2-a multiple sequence alignment editor and analysis workbench. Bioinformatics 25, 1189–1191.

Wu, H., Zhu, H., Miao, H., Perelson, A.S., 2008. Parameter identifiability and estimation of HIV/AIDS dynamic models. Bulletin of Mathematical Biology 70, 785–799.

Yang, Z., 1996. Among-site rate variation and its impact on phylogenetic analyses. Trends in Ecology & Evolution 11, 367–372.

Yang, Z., Rannala, B., 1997. Bayesian phylogenetic inference using DNA sequences: a Markov Chain Monte Carlo method. Molecular Biology and Evolution 14, 717–724.

Yildirim, S., Singh, S.S., Dean, T., Jasra, A., 2015. Parameter estimation in hidden markov models with intractable likelihoods using sequential monte carlo. Journal of Computational and Graphical Statistics 24, 846–865.

